# Genetic analyses in UK Biobank identifies 78 novel loci associated with urinary biomarkers providing new insights into the biology of kidney function and chronic disease

**DOI:** 10.1101/315259

**Authors:** Daniela Zanetti, Abhiram Rao, Stefan Gustafsson, Themistocles Assimes, Stephen B. Montgomery, Erik Ingelsson

**Author notes:** **Address for Correspondence**: Erik Ingelsson, MD, PhD, FAHA, 300 Pasteur Dr, mail code: 5773; Stanford, CA 94305; USA.

## Abstract

**Background:** Urine biomarkers, such as creatinine, microalbumin, potassium and sodium are strongly associated with several common diseases including chronic kidney disease, cardiovascular disease and diabetes mellitus. Knowledge about the genetic determinants of the levels of these biomarker may shed light on pathophysiological mechanisms underlying the development of these diseases.

**Methods:** We performed genome-wide association studies of urinary levels of creatinine, microalbumin, potassium, and sodium in up to 326,441 unrelated individuals of European ancestry from the UK Biobank, a large population-based cohort study of over 500,000 individuals recruited across the United Kingdom in 2006-2010. Further, we explored genetic correlations, tissue-specific gene expression and possible causal genes related to these biomarkers.

**Results:** We identified 23 genome-wide significant independent loci associated with creatinine, 20 for microalbumin, 12 for potassium, and 38 for sodium. We confirmed several established associations including between the *CUBN* locus and microalbumin (rs141640975, p=3.11e-68). Variants associated with the levels of urinary creatinine, potassium, and sodium mapped to loci previously associated with obesity (*GIPR,* rs1800437, p=9.81e-10), caffeine metabolism (*CYP1A1*, rs2472297, p=1.61e-8) and triglycerides (*GCKR,* rs1260326, p=4.37e-16), respectively. We detected high pairwise genetic correlation between the levels of four urinary biomarkers, and significant genetic correlation between their levels and several anthropometric, cardiovascular, glycemic, lipid and kidney traits. We highlight *GATM* as causally implicated in the genetic control of urine creatinine, and *GIPR*, a potential diabetes drug target, as a plausible causal gene involved in regulation of urine creatinine and sodium.

**Conclusion:** We report 78 novel genome-wide significant associations with urinary levels of creatinine, microalbumin, potassium and sodium in the UK Biobank, confirming several previously established associations and providing new insights into the genetic basis of these traits and their connection to chronic diseases.

**Author Summary:** Urine biomarkers, such as creatinine, microalbumin, potassium and sodium are strongly associated with several common diseases including chronic kidney disease, cardiovascular disease and diabetes mellitus. Knowledge about the genetic determinants of the levels of these biomarker may shed light on pathophysiological mechanisms underlying the development of these diseases. Here, we performed genome-wide association studies of urinary levels of creatinine, microalbumin, potassium and sodium in up to 326,441 unrelated individuals of European ancestry from the UK Biobank. Further, we explored genetic correlations, tissue-specific gene expression and possible causal genes related to these biomarkers. We identified 78 novel genome-wide significant associations with urinary biomarkers, confirming several previously established associations and providing new insights into the genetic basis of these traits and their connection to chronic diseases. Further, we highlight *GATM* as causally implicated in the genetic control of urine creatinine, and *GIPR*, a potential diabetes drug target, as a plausible causal gene involved in regulation of urine creatinine and sodium. The knowledge arising from our work may improve the predictive utility of the respective biomarker and point to new therapeutic strategies to prevent common diseases.

## Introduction

Fluctuating levels of several urinary biomarkers are used clinically to assess an individual’s renal function as well as to diagnose and predict the onset of related chronic diseases(1). These biomarkers include creatinine, microalbumin, potassium, sodium, and other proteins and peptides which have been associated with chronic kidney disease (CKD) (2, 3) cardiovascular disease (CVD) (4-8), and type 2 diabetes (T2D) (9, 10). Urinary biomarkers have also shown promise in monitoring response to therapy (11). In comparison to blood, biomarkers in urine are less subject to homeostatic mechanisms. This situation allows for greater fluctuations of biomarker levels which in turn may provide a signal that more reliably reflects dynamic changes in human biological and pathophysiological processes (12).

Little progress has been made in disentangling the genetic determinants of levels of urinary biomarkers in large population cohorts despite extensive research on the genetic determinants of biomarkers in blood (13-16), including estimated glomerular filtration rate (17). Discovering such associations and identifying whether they are genetically correlated with other common traits and physiological metrics may provide important etiological insights into their control. This knowledge may in turn improve the predictive utility of the respective biomarker and point to new therapeutic strategies to prevent common diseases.

In this context, we performed a genome-wide association study (GWAS) of four urine biomarkers individually - creatinine, microalbumin, potassium and sodium - in up to 326,441 participants of the UK Biobank (UKB) study. For each urinary biomarker, we estimated its heritability, identified genetic associations that were likely mediated by expression or methylation quantitative trait loci (eQTLs or mQTLs), evaluated genetic correlations with several anthropometric, cardiovascular, glycemic, lipid, hematological and kidney traits, and used a bioinformatics approach to pinpoint tissues that were significantly enriched for associated variants, as well as candidate causal genes.

## Results

### Association analyses

We found a total of 93 genome-wide significant independent variants associated with any of the four urine biomarkers: 23 for creatinine (N = 327,857), 20 for microalbumin (N = 326,441), 12 for potassium (N = 327,147), and 38 for sodium (N = 327,162) (Table 1, Figures 1a-d). A total of 85 lead variants are novel, while 8 are located in loci previously associated with kidney function, including *CUBN (35)*, *CPS1*, *GATM* and *SHROOM3 (36)*. Out of the 85 novel variant associations, three were shared across creatinine, potassium and sodium (rs2472297, rs4410790 and rs784257) and one between creatinine and potassium (rs13143189); consequently, we report a total of 78 novel and unique loci associated with urinary biomarkers.

**Figure 1.**
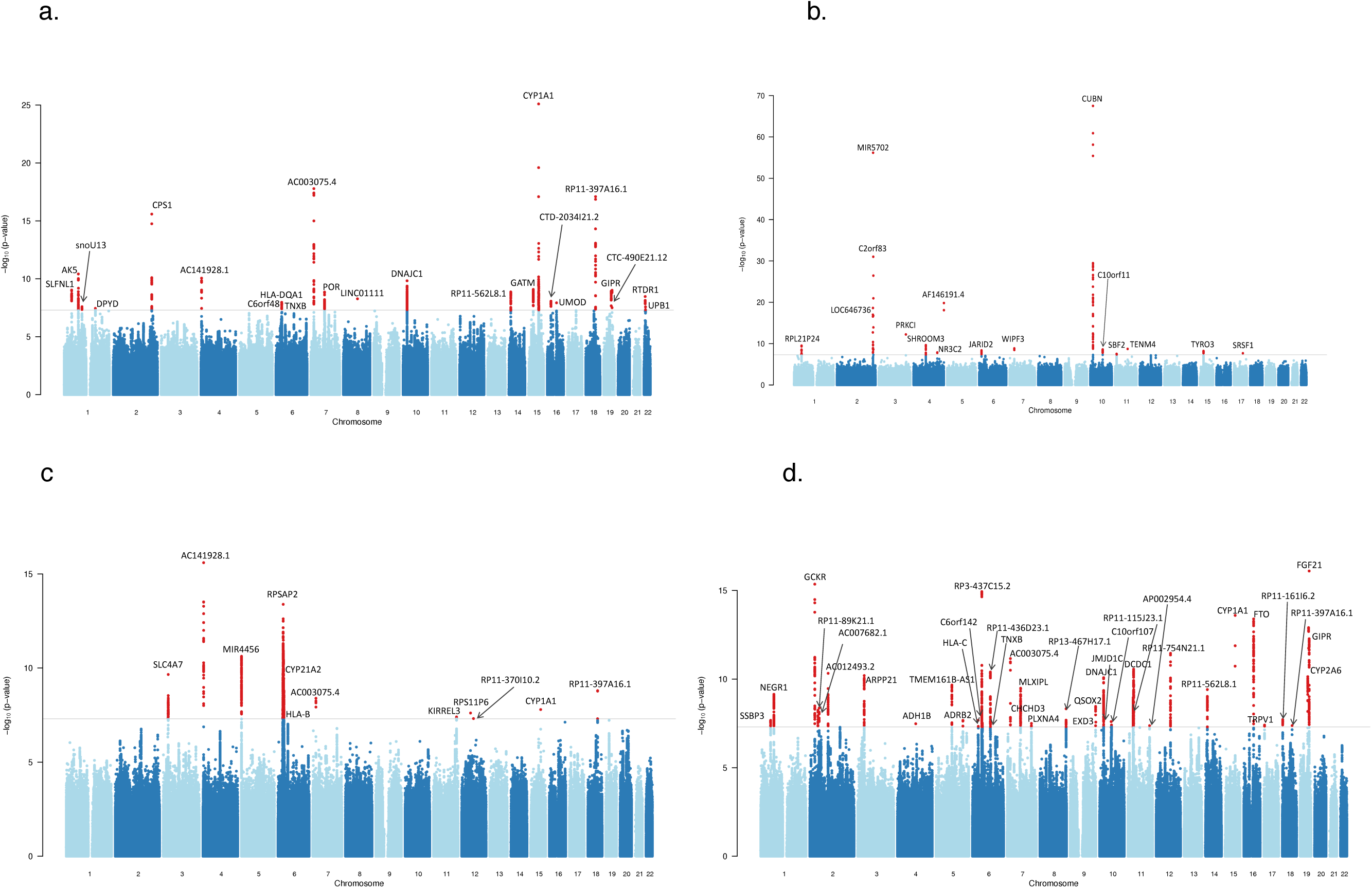
Manhattan plot for genetic associations for urinary creatinine (a), microalbumin (b), potassium (c) and sodium (d). The nearest gene for each chromosome labeled. Negative log_10_-transformed *P* values for each SNP (*y* axis) are plotted by chromosomal position (*x* axis). The gray line represents the threshold for genome-wide statistically significant associations (*P* = 5 × 10^−8^). Red points represent significant hits.

**Table 1.**
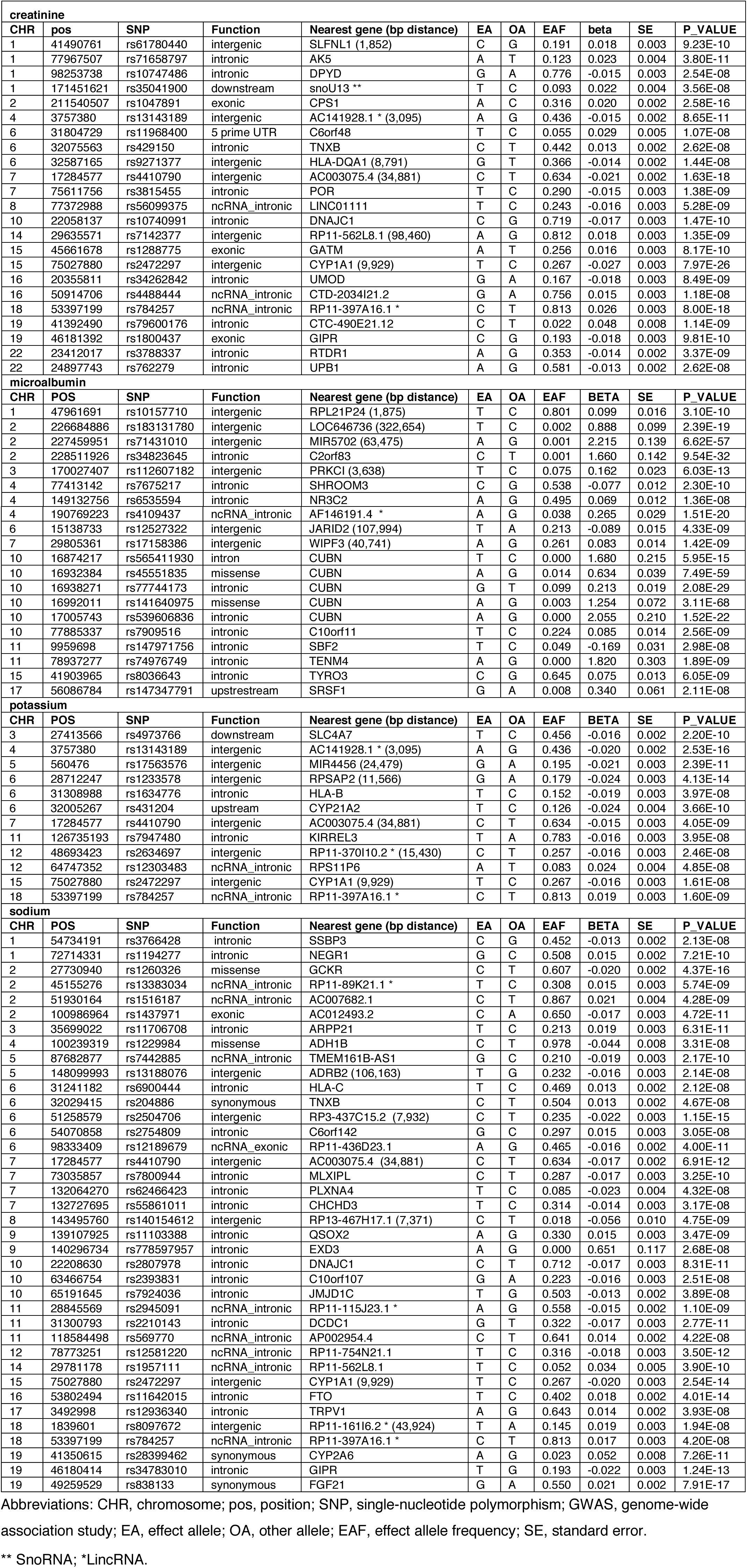
Genetic loci associated with urine biomarkers in the genome-wide association studies.

Many of the 78 novel lead SNPs associated with any of the four urine biomarkers are located near loci (± 250 kb) previously reported to be associated with several complex diseases/traits as compiled by the GWAS catalog (S2 Table). For our creatinine loci, the strongest associations (based on the lowest p-values in p-value) were found for height (37), lung cancer (38), schizophrenia (39), human blood cells (40), coffee consumption (41), basal cell carcinoma (42), breast cancer (43), BMI (44), kidney function (17), Crohn’s disease (45), intraocular pressure (46), circulating parathyroid hormone (47), and liver enzyme in plasma (48). Only one locus (rs13143189) had no associations with GWAS catalog traits. For our 14 microalbumin loci, the strongest associations were found with male-pattern baldness (49), educational attainment (50), human blood cells (40), allergic disease (51), and prostate cancer (52). Furthermore, 5 variants are located near *CUBN*, a well-known locus for albuminuria (35) (Table 1) and 6 loci have no previously known associations with traits in the GWAS catalog. For potassium, the strongest associations were found with breast cancer (53), esophageal adenocarcinoma (54), schizophrenia (39), human blood cells (40), neuroticism (55), and resting heart rate (56) (S2 Table). For sodium, the strongest associations were found with human blood cells (40), BMI (57), lipid levels (58), fasting plasma glucose (59), educational attainment (50), intelligence (60), neuroticism (55), alcohol consumption (61), retinal vascular caliber (62), liver enzyme in plasma (48), triglycerides (63), schizophrenia (39), breast cancer (53) blood pressure (64), non-glioblastoma glioma (65), lung cancer (38), and mumps (66). A total of 7 loci had no previously known association with a GWAS catalog trait.

Regional plots for the 92 discovered loci are shown in S1-S4 Figs. We observed minor inflation in test statistics (lambda=1.149 for creatinine, 1.033 for microalbumin, 1.107 for potassium, and 1.158 for sodium), which is expected (67) under polygenic inheritance in large samples (S5a-d Fig). The gene-level association identified 51, 11, 31 and 42 genes associated at GWAS significance with creatinine, microalbumin, potassium and sodium, respectively (S6 Fig, S3 Table). Several of these genes have been previously associated with complex traits, such as *CPS1*, *GATM/SPATA5L1* and *GIPR* (creatinine) which have been implicated in the development of atherosclerosis (68), CKD (17) (36) and obesity (44), respectively. We confirmed the association with *CUBN* (microalbumin), a well-known locus for albuminuria (35). The potassium locus including *ELL* showed association with several cancers including prostate (18), lung (69) and esophagus (70), while the *FTO* locus for sodium coincides with the first and strongest locus identified to date for BMI through GWAS (44).

### LD score regression

We found evidence of high genetic correlation between every pair of the 4 urinary biomarkers, although the correlations between microalbumin and the other three biomarkers were generally lower (0.20-0.28), than between the other three (0.53-0.81). Further, we observed significant genetic correlation between the urinary biomarkers and several anthropometric, cardiovascular, glycemic, lipid, hematologic and kidney traits (Figure 2, S4 Table). Specifically, we identified a significant and positive genetic correlation across creatinine, microalbumin and sodium and several traits related to cardiometabolic disease including BMI, body fat, obesity, WHR, fasting insulin, and triglycerides. In addition, we detected significant negative correlation between urinary creatinine and eGFRcrea; and between three urinary biomarkers (creatinine, microalbumin and sodium) and HDL. We observed a significant and positive correlation between potassium and obesity, and a strong inverse correlation between potassium and systolic blood pressure. The heritability of the four urine biomarkers was in the range of 0.015 ≤ h^2^_g_ ≤ 0.069 (Table 2).

**Figure 2.**
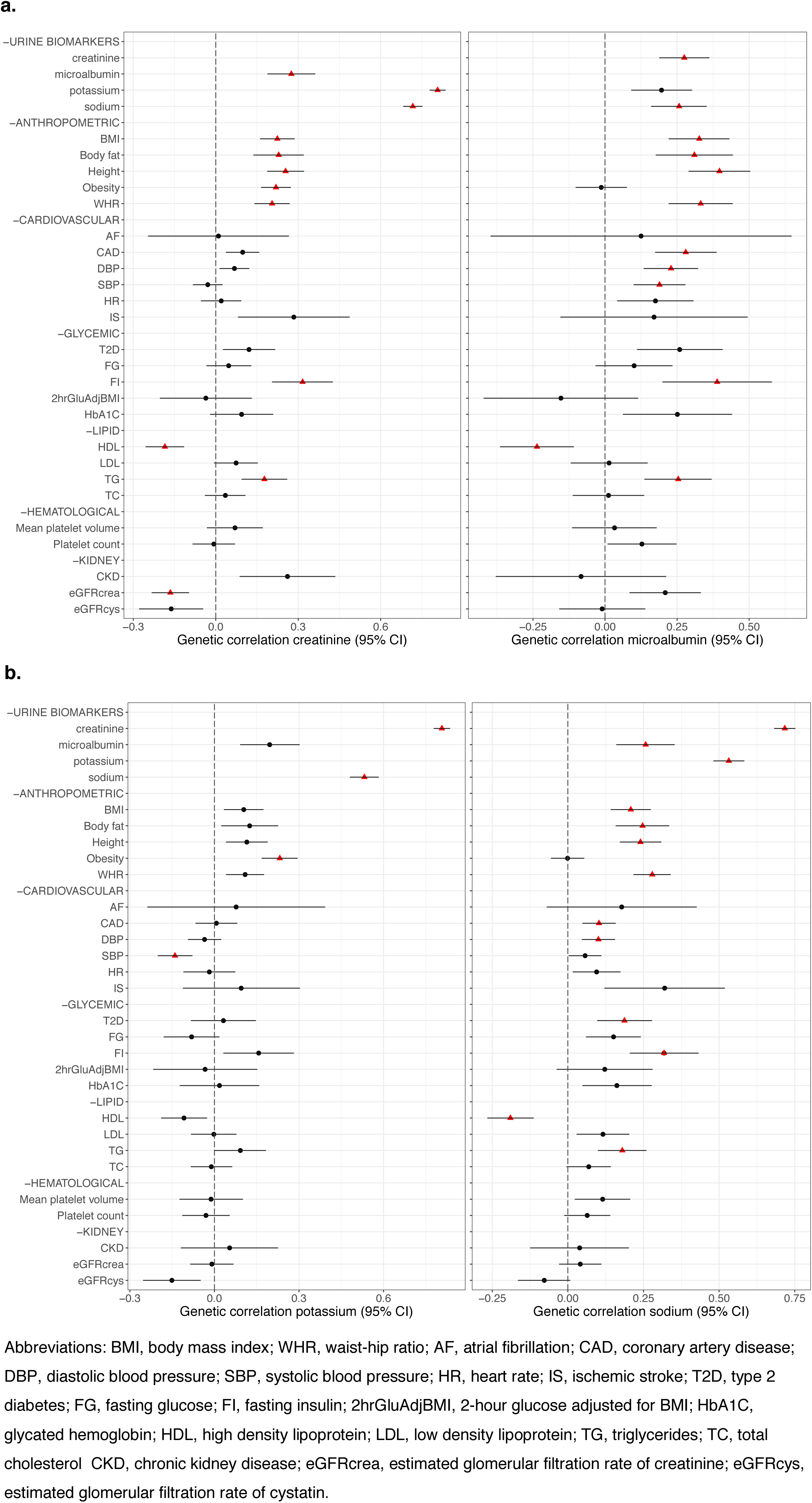
Genetic correlations between urinary creatinine, microalbumin (a), potassium and sodium (b) and other traits. Significant correlations after Bonferroni correction (4.46 × 10^−4^) are highlighted with a red triangle.

**Table 2.**
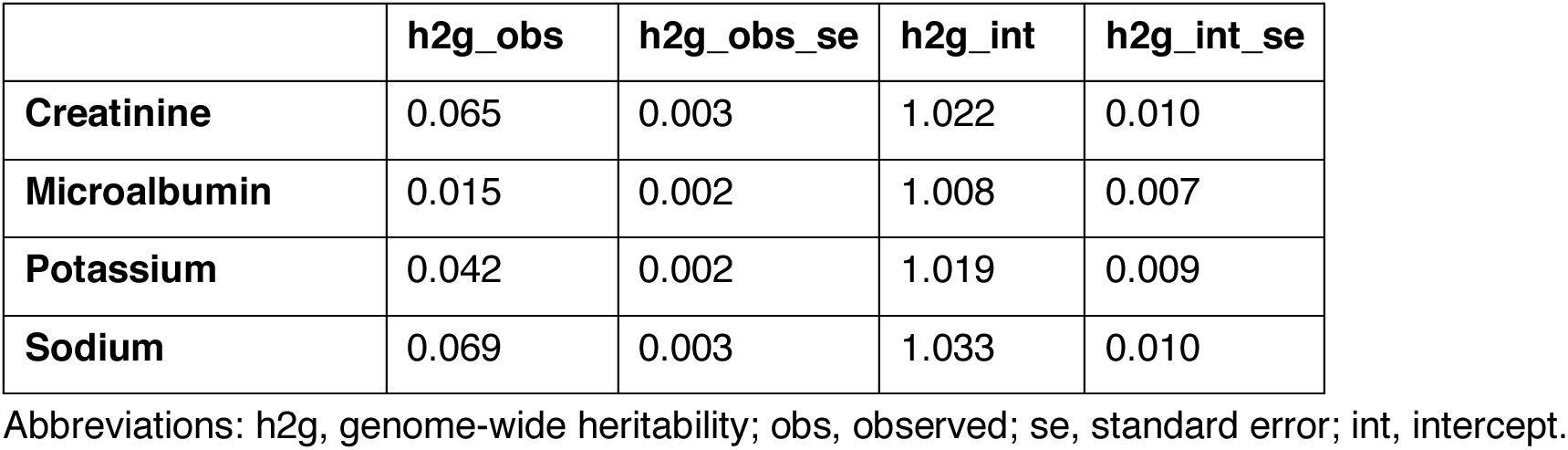
Genome-wide heritability of the four urine biomarkers.

|  | **h2g_obs** | **h2g_obs_se** | **h2g_int** | **h2g_int_se** |
| --- | --- | --- | --- | --- |
| **Creatinine** | 0.065 | 0.003 | 1.022 | 0.010 |
| **Microalbumin** | 0.015 | 0.002 | 1.008 | 0.007 |
| **Potassium** | 0.042 | 0.002 | 1.019 | 0.009 |
| **Sodium** | 0.069 | 0.003 | 1.033 | 0.010 |

Abbreviations: h2g, genome-wide heritability; obs, observed; se, standard error; int, intercept.

### Functional analysis

Most of the significant SNP associations we identified were located in intronic or intergenic regions, rather than in exons or regulatory regions (S7-8 Figs). Most genome-wide significant SNPs across the four urine biomarkers were predicted to have no or weak effects on transcription based on the prevalent minimum (most active) chromatin state (S9 Fig). DEPICT analyses identified hematologic and immune systems as the most enriched physiological systems for loci associated with creatinine; cardiovascular systems for microalbumin; and nervous systems for potassium and sodium biomarkers. However, none of the DEPICT analyses reached statistical significance (FDR<0.05, S10 Fig, S5 Table). Genes identified by gene-level associations as associated with creatinine were enriched among upregulated DEG sets in the liver, and genes associated with microalbumin were enriched among downregulated gene sets in the bladder (Bonferroni corrected P-value ≤ 0.05). The other biomarkers did not show any significant relationship across the genes expressed in the 53 tissue types of GTEx (S11-12 Fig).

### Colocalization with eQTL and mQTL summary statistics

We detected four significant eQTL probe colocalizations for creatinine (S6 Table). No significant eQTL colocalization signals were detected for microalbumin, potassium and sodium. We observed 43 significant mQTL probe colocalizations for creatinine, two for microalbumin, 157 for potassium and 30 for sodium; however, many probes were found to map to the same region and are likely representing the same methylation effect (S7 Table). We observed a total of 24 independent (r^2^ for top SNPs > 0.5) and high confidence colocalizations at loci with significant eQTL and/or mQTL SMR p-values (Table 3).

**Table 3.** Consistent significant signals across genome-wide association studies, expression quantitative trait loci (eQTL) and/or methylation quantitative trait loci (mQTL) probe colocalizations.

| eQTL |  |  |  |  |  |  |  |  |  |  |  |  |  |  |
| --- | --- | --- | --- | --- | --- | --- | --- | --- | --- | --- | --- | --- | --- | --- |
|  | ProbeChr | topSNP_bp | topSNP | Nearest gene | A1 | A2 | Freq | b_GWAS | se_GWAS | p_GWAS | b_eQTL | se_eQTL | p_eQTL | p_SMR |
| creatinine | 7 | 75657850 | rs11983987 | POR | G | A | 0.182 | -0.017 | 0.003 | 2.85E-08 | 0.4 | 0.026 | 1.41E-54 | 1.72E-07 |
|  | 15 | 45711492 | rs2467858 | GATM | G | A | 0.276 | 0.016 | 0.003 | 1.31E-09 | -0.383 | 0.022 | 2.44E-70 | 9.51E-09 |
|  | 15 | 45685487 | rs9788780 | SPATA5L1 | A | T | 0.273 | 0.016 | 0.003 | 1.19E-09 | 0.85 | 0.019 | 8.63E-103 | 1.67E-09 |
|  | 15 | 75622943 | rs9673084 | COMMD4 | A | G | 0.245 | -0.016 | 0.003 | 2.92E-10 | 0.139 | 0.022 | 1.48E-10 | 7.01E-06 |
| mQTL |  |  |  |  |  |  |  |  |  |  |  |  |  |  |
| creatinine | 1 | 41487317 | 41560291 | SCMH1 | A | G | 0.197 | 0.016 | 0.003 | 6.20E-09 | 0.629 | 0.037 | 3.40E-64 | 3.88E-08 |
|  | 1 | 78442554 | 78623626 | GIPC2,LOC100132264 | T | C | 0.1 | 0.021 | 0.003 | 6.44E-10 | 0.527 | 0.046 | 4.99E-30 | 5.61E-08 |
|  | 4 | 3748154 | 3748134 | LOC100129786,ADRA2C | A | C | 0.404 | 0.015 | 0.002 | 9.28E-10 | -0.636 | 0.032 | 2.61E-88 | 4.87E-09 |
|  | 6 | 32165183 | 32587165 | HLA-DRB1,HLA-DQA1 | G | T | 0.383 | -0.014 | 0.002 | 1.44E-08 | -0.322 | 0.034 | 2.45E-21 | 1.14E-06 |
|  | 7 | 17337976 | 17338147 | LOC100131512 | T | C | 0.136 | 0.019 | 0.003 | 1.46E-08 | 0.419 | 0.047 | 3.84E-19 | 1.70E-06 |
|  | 7 | 75624427 | 75695081 | MDH2 | A | G | 0.173 | -0.017 | 0.003 | 1.74E-08 | -0.69 | 0.042 | 1.72E-61 | 9.55E-08 |
|  | 15 | 74891207 | 75027880 | CYP1A1,CYP1A2 | T | C | 0.231 | -0.027 | 0.003 | 7.97E-26 | 0.264 | 0.037 | 9.21E-13 | 3.49E-09 |
|  | 15 | 75165896 | 75328595 | SCAMP2 | C | T | 0.335 | -0.016 | 0.002 | 1.61E-10 | -0.531 | 0.032 | 4.05E-61 | 2.49E-09 |
|  | 19 | 46181546 | 46182304 | GIPR | A | T | 0.208 | -0.018 | 0.003 | 1.52E-09 | -0.804 | 0.039 | 2.13E-93 | 6.81E-09 |
| microalbumin | 4 | 77409957 | 77413179 | SHROOM3 | G | A | 0.489 | 0.077 | 0.012 | 2.83E-10 | 0.268 | 0.033 | 0 | 6.99E-07 |
|  | 11 | 10323902 | 10269178 | ADM | G | A | 0.22 | 0.079 | 0.015 | 2.36E-07 | -0.63 | 0.041 | 0 | 9.52E-07 |
| potassium | 4 | 3748154 | 3748134 | LOC100129786,ADRA2C | A | C | 0.404 | 0.018 | 0.003 | 3.95E-12 | -0.636 | 0.032 | 2.61E-88 | 5.64E-11 |
|  | 5 | 601475 | 599269 |  | A | G | 0.198 | -0.018 | 0.003 | 2.23E-09 | -0.991 | 0.035 | 4.23E-177 | 4.87E-09 |
|  | 6 | 28830902 | 28934352 |  | C | T | 0.073 | -0.025 | 0.004 | 2.44E-11 | -1.597 | 0.039 | 8.63E-120 | 4.35E-11 |
|  | 15 | 74891207 | 75027880 | CYP1A1,CYP1A2 | T | C | 0.231 | -0.016 | 0.003 | 1.61E-08 | 0.264 | 0.037 | 9.21E-13 | 9.38E-06 |
| sodium | 7 | 72993570 | 72991704 | TBL2 | A | G | 0.269 | -0.015 | 0.003 | 3.94E-09 | 0.755 | 0.034 | 1.68E-109 | 1.27E-08 |
|  | 8 | 143484414 | 143527647 |  | T | C | 0.016 | -0.043 | 0.008 | 2.65E-08 | -1.178 | 0.11 | 5.79E-27 | 7.77E-07 |
|  | 15 | 74891207 | 75027880 | CYP1A1,CYP1A2 | T | C | 0.231 | -0.02 | 0.003 | 2.54E-14 | 0.264 | 0.037 | 9.21E-13 | 1.88E-07 |
|  | 19 | 46181546 | 46182304 | GIPR | A | T | 0.208 | -0.022 | 0.003 | 1.87E-13 | -0.804 | 0.039 | 2.13E-93 | 4.34E-12 |
|  | 19 | 49259452 | 49259529 | FGF21 | A | G | 0.418 | -0.021 | 0.002 | 7.91E-17 | 0.386 | 0.032 | 7.59E-33 | 8.32E-12 |

Abbreviations: CHR chromosome; pos, position; SNP, single-nucleotide polymorphism; EA, effect allele; OA, other allele; EAF, effect allele frequency; b, beta; p, p-value; SMR, summary-based-results Mendelian Randomization.

#### • Creatinine

We detected significant eQTL colocalizations for *GATM* (lead SNP rs2467858, SMR p = 9.51e-9) and *SPATA5L1* (lead SNP rs9788780, SMR p = 1.67e-9) on chromosome 15 (Figure 3,Table 3); these GWAS signals colocalized with several mQTL probes in the region as well (S7 Table). The most strongly associated GWAS variant was rs1288775 (beta = 0.016, p-value = 8.17e-10) (Table 1), and all colocalization lead variants at this locus were strongly linked (LD r^2^ > 0.9). We identified an eQTL colocalization on chromosome 15 at the *COMMD4* locus (lead SNP rs9673084, p = 7.01e-6). We also observed two additional clusters of mQTL colocalizations on chromosome 15, one set with lead variants including eQTLs (or linked variants) for *CYP1A1*/*CYP1A2* (top probe SNP rs2472297, p = 3.49e-9), and another set with lead variants including eQTLs for *SCAMP2* (top probe lead SNP rs4886649, p = 2.49e-9). However, we did not observe eQTL colcalizations for both these genes in blood. We also observed an eQTL colocalization at the *POR* locus on chromosome 7 (lead SNP rs11983987, p=1.72e-7). Although there were no mQTL colocalizations with lead SNPs in strong linkage with rs11983987, we detected 4 colocalizations of mQTL probes with lead SNPs in weak linkage (LD r2 ~ 0.4). The SNPs rs13143189 (p = 8.56e-11), rs9271377 (p = 1.44e-8), rs2472297 (p = 7.97e-26) and rs1800437 (p = 9.81e-10) showed significant association in the GWAS analysis and significant colocalization with methylation probes. These SNPs are close to *ADRA2C* on chromosome 4, *HLA-DQA1* on chromosome 6, *CYP1A2* on chromosome 15 and *GIPR* on chromosome 19, respectively (Table 3).

**Figure 3.**
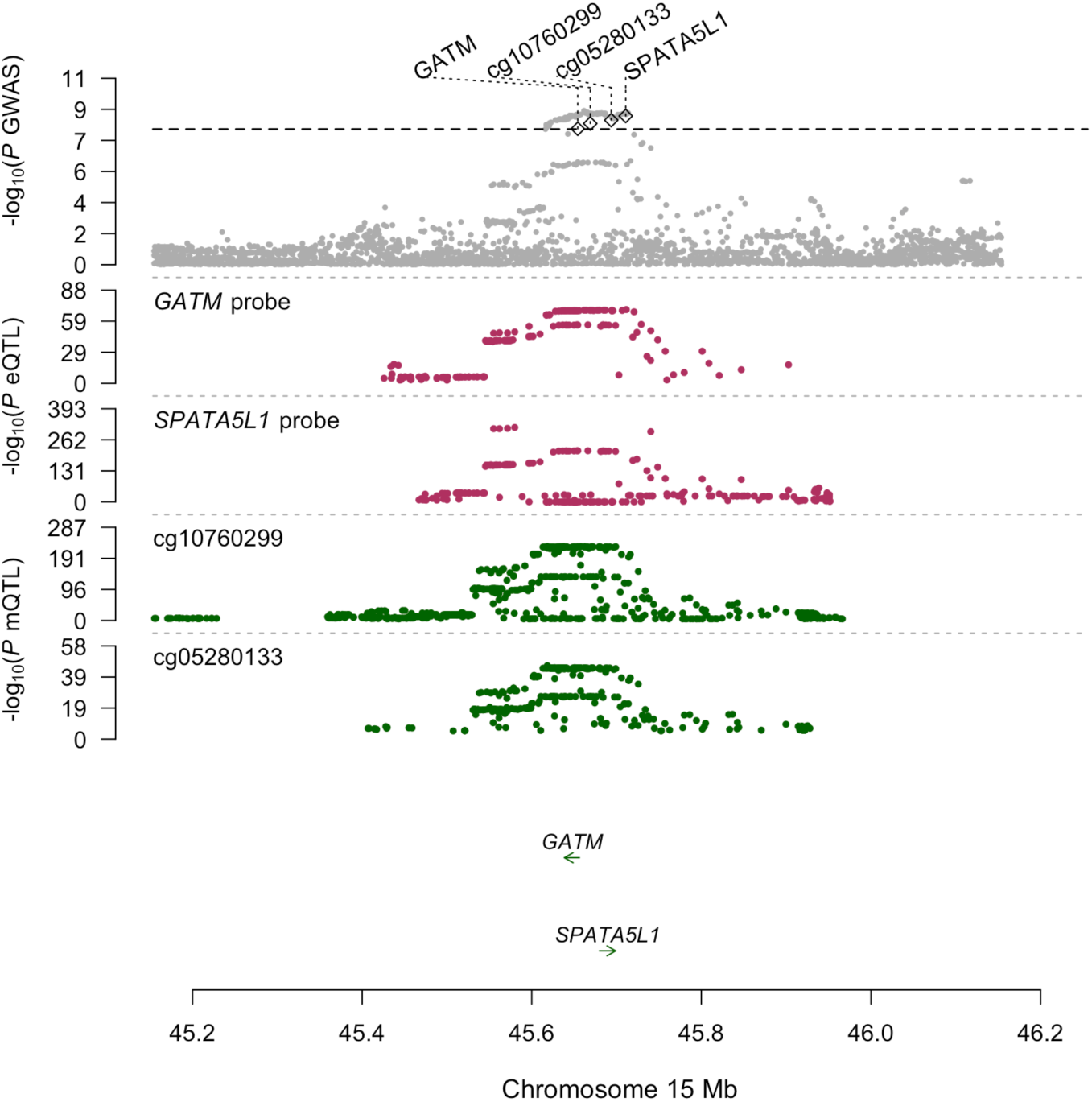
Expression quantitative trait loci (eQTL) and methylation quantitative trait loci (mQTL) colocalization probes for the locus *GATM*/*SPATA5L1*. Colocalization of GWAS, eQTL and mQTL signals at the *GATM* locus on chromosome 15. eQTL signals for *GATM* and *SPATA5L1*, and the two top mQTL probes are shown, along with their respective colocalization top variants (determined by maximum SMR effect size)

#### • Microalbumin

We detected significant mQTL colocalizations for *ADM* on chromosome 11 (lead SNP rs4910113, SMR p = 9.52e-7) and consistent signals in GWAS and mQTL colocalization analyses for *SHROOM3* on chromosome 4 (rs7675258, p = 6.99e-7).

#### • Potassium

We detected consistent signals in GWAS and mQTL colocalization analyses for the *CYP1A1* locus on chromosome 15 (rs2472297, p = 9.38e-6). Most mQTL probe colocalizations (n = 134) for potassium occurred for probes binding within a 5.5 Mb region on chromosome 6 (chr6:27277996-32729059), which overlaps with the HLA region. These colocalizations clustered intro three broad groups based on linkage of their lead variants; identification of genes is complicated by the linkage structure of this region of the genome (S7 Table).

#### • Sodium

We detected consistent significant mQTL signals for the *TBL2* locus on chromosome 7 (rs12540011, p = 1.27e-8), for the *CYP1A1* locus on chromosome 15 (rs2472297, p = 1.88e-7), and for the *GIPR* (rs10423928, p = 4.34-12) and *FGF21* loci (rs838133, p = 8.32e-12) on chromosome 19.

We also detected shared mQTL colocalizations between urine biomarkers. Creatinine and potassium shared an mQTL colocalization (with multiple probes) on chromosome 4; the lead SNP rs12641452 of the top probe is located within a *POLR2A* binding site in a 25kb region flanking *ADRA2C*. Creatinine and sodium shared an mQTL colocalization near *GIPR* on chromosome 19 with the same lead variant rs10423928 identified across multiple probes. The methylation probes that colocalize with the GWAS signal at this locus bind in or near a high confidence *CTCF* binding site in an intronic region of *GIPR*.

## Discussion

### Principal findings

We achieved three broad goals in this study of up to 326,441 unrelated UKB participants of European descent. We (1) established the genetic determinants of levels of creatinine, microalbumin, potassium and sodium in urine, (2) evaluated genetic correlation between these biomarkers and several physiological measurements, and (3) explored the functional impact of associated loci using variant annotations, tissue-specific gene expression patterns, and colocalization analysis with expression and methylation summary statistics. Our main findings are several-fold. First, we report a total of 78 novel loci associated with one or more of the 4 urine biomarkers, providing new leads regarding biological processes involved in regulating urinary creatinine, microalbumin, potassium and sodium. Second, our analyses indicate low heritability, but high pairwise genetic correlation for the four urinary biomarkers; as well as significant genetic correlations with several traits related to chronic kidney disease, cardiovascular disease, and type 2 diabetes. Third, we highlight a significant positive relationship for creatinine and microalbumin and genes highly expressed in liver and bladder, respectively. Fourth, we identify 4 and 20 independent colocalization events of GWAS data with blood gene expression and DNA methylation respectively, and we provide evidence for a functional mechanism at colocalized GWAS loci. As an example of the latter, we identified *GATM* and *SPATA5L1* as possible causal genes involved in the genetic underpinning of urinary creatinine; and *GIPR*, a potential drug target in the treatment of obesity-associated metabolic disorders, as a plausible causal gene involved in regulation of urine creatinine and sodium.

### Comparison with prior literature

This study is the first GWAS of urinary creatinine, potassium and sodium to the best of our knowledge while genetic determinants of microalbumin have already been explored in previous, smaller studies (10, 35). In this study, we replicated the well-known association in *CUBN*, and extend prior GWAS of microalbumin by highlighting several novel pathways influencing this trait. We also replicated the known association of variants in the *CPS1* locus with serum creatinine, previously reported to affect creatinine production and secretion; and with *GATM,* a gene that encodes a mitochondrial enzyme previously associated with serum creatinine (36).

Previous epidemiological studies have showed significant associations between sodium and potassium intake and CVD (6, 7), and between higher levels of sodium and lower levels of potassium intake and higher blood pressure (8), but there is a lack of studies studying the genetics underpinning of these biomarkers. Our genetic correlation analysis confirmed a negative association of blood pressure with urinary potassium, and the positive association with urinary sodium. We also confirmed a significant positive genetic correlation between creatinine, microalbumin and sodium and several anthropometric measurements related with CVD (BMI, body fat, obesity and WHR), and with fasting insulin, and triglycerides. In addition, we detected significant negative correlations of urinary creatinine, microalbumin and sodium with HDL.

We report heritability estimates of the 4 urinary biomarkers in the range of 2-7%, which is lower than many complex traits, but quite consistent with prior studies of kidney-related biomarkers. Previous studies applying family-based designs have generally reported similar, but somewhat higher estimates for other kidney-related biomarkers (71-73). Specifically, the estimates for albumin/creatinine ratio, potassium and sodium have been reported to be in the range of 13-19 %, lowest for potassium and highest for creatinine (71, 73). This discrepancy can probably be explained by prior studies focusing on serum biomarkers, and due to differences in statistical methods used. Indeed, our method captures SNP-based heritability due to common variation, whereas twin and other family-based analyses also capture components of heritability due to rare variation (74).

### Novel biology

We observed colocalization of the GWAS signal for urine creatinine at *GATM* and *SPATA5L1* with both eQTL and mQTL signals in the region. The *GATM* locus is not novel in terms of association with renal function as previous studies have shown its association with CKD and reduced glomerular filtration rate (17, 36). *GATM,* which stands for glycine amidinotransferase, encodes a rate-limiting enzyme involved in creatinine biosynthesis, and has been suggested to act as a functional link between statin-mediated lowering of cholesterol and susceptibility to statin-induced myopathy (75). In addition, a previous study showed a significant association of SNPs in the *GATM* locus with plasma and urine creatinine but not with cystatin C in plasma, another biomarker of renal function (36). This association may indicate that the regulatory variant(s) in this locus influence creatinine production rather than creatinine excretion. Consistent with this finding, our genetic correlation analysis indicated a significant negative association of urine and eGFRcrea (p = 2.08e-06), but no significant correlation with eGFRcys, pointing more broadly to common pathways between urine and serum creatinine. The colocalization lead variants for all eQTL and mQTL probes in the region were strongly linked with each other and with the previously reported statin-induced response eQTL rs9806699 for *GATM* (LD r^2^ > 0.9). Consistent with the expectation that increased methylation of CpG-dense promoters correlates negatively with gene expression, the linked lead eQTL and mQTL colocalization SNPs showed association with decreased *GATM* expression and increased methylation in the region (S6 Table).

Variants in *SHROOM3* showed significant association with microalbumin in the GWAS and in the mQTL colocalization. *SHROOM3* encodes an actin-associated protein important in epithelial morphogenesis that has been previously shown to be strongly associated with kidney function (36). Studies in zebrafish and rat show that alterations in *SHROOM3* can result in glomerular dysfunction. Furthermore, human *SHROOM3* variants can induce impaired kidney function in animal models (76). Variants near *CYP1A1* on chromosome 15 showed a significant association with urinary creatinine, potassium and sodium in the GWAS; and in the mQTL colocalization analysis. This locus has been previously suggested as a potential drug target for the prevention of CVD because variants in this locus are also associated with blood pressure (77). Likewise, variants in the *GIPR* locus significantly associated with creatinine and sodium in our GWAS and mQTL colocalization have been previously associated with several cardiometabolic traits including obesity (78), BMI (44), and hip circumference (79). The glucose-dependent insulinotropic peptide (GIP) has a central role in glucose homeostasis through its amplification of insulin secretion. Incretin hormones such as GIP act to promote efficient uptake and storage of energy after food ingestion and have become important players for glucose homeostasis in pancreatic and extra pancreatic tissue. A recent study demonstrated that mice with selective ablation of GIPR in beta cells exhibited lower levels of meal-stimulated insulin secretion, decreased expansion of adipose tissue mass and preservation of insulin sensitivity when compared to controls (80). Hence, the GIPR represents a potential therapeutic in the treatment of diabetes. Similarly, the fibroblast growth factor 21 (*FGF21*) significant associated with sodium in our GWAS and mQTL colocalization, is considered a novel promising candidate in the treatment of T2D, obesity, dyslipidemia, cardiovascular and fatty liver diseases (81). *FGF21* encodes a fibroblast growth factor involved in glucose and lipid metabolism, and has been previously associated with macronutrient intake (82). In vitro, *FGF21* promotes insulin-dependent glucose uptake through the transcription of GLUT1 in rodent and human adipocytes (83). Pharmacologic doses of *FGF21* improve glucose clearance and insulin sensitivity, and lower plasma triglycerides and free fatty acids in diabetic and obese animal models (84, 85). In humans, the role of *FGF21* remains ill-defined, but reports have linked serum levels of *FGF21* with adiposity, fasting insulin, and triglycerides (86).

### Strengths and limitations

Strengths of the present study include the very large sample size with both genetic profiling and phenotypic data which enabled us to detect a large number of genetic associations, the use of state-of-the-art methods to validate our results including a conservative analytical framework with strict multiple testing correction, and a variety of pathway analyses. Furthermore, our study is the most comprehensive to date on the genetics of urine biomarkers, combining GWAS, genetic correlation, as well as functional and eQTL and mQTL colocalization analyses.

We also acknowledge some limitations. First, we did not replicate our findings in an external study sample due to unavailability of a similar independent large study; although we applied an internal replication strategy given the very large sample size. Second, participants included in our analyses were restricted to middle-aged and elderly individuals of European ancestry potentially limiting the generalizability of our results to other age groups and ethnicities.

### Conclusions

We report 78 novel genome-wide significant associations with urinary creatinine, microalbumin, potassium and sodium in the UKB, confirming several known associations and providing new insights into the genetic basis of these traits and their connection to chronic diseases. We detected high genetic correlations between the four urinary biomarkers and significant genetic correlations with several anthropometric, cardiovascular, glycemic, lipid and kidney traits. Through this effort, we highlight *GATM* as a plausible causal gene controlling levels of urinary creatinine, and *GIPR* – a potential diabetes drug target – as being associated with urine creatinine and sodium.

## Materials and Methods

### Study sample

The UKB is a longitudinal cohort study of over 500,000 individuals aged 40-69 years at the time of recruitment between 2006 and 2010. Participants were enrolled in 22 study centers located in England, Scotland and Wales. The UKB study was approved by the North West Multi-Centre Research Ethics Committee and all participants provided written informed consent. Details of these measurements can be found in the study protocol (18) and in the UKB Data Showcase (http://biobank.ctsu.ox.ac.uk/crystal/).

### Phenotype

Urine samples were collected at baseline in all UKB participants. All urinary biomarker measurements were carried out on a single Beckman Coulter AU5400 clinical chemistry analyzer using the manufacturer’s reagents and calibrators, except for urinary microalbumin, which used reagents and calibrators sourced from Randox Bioscience. Internal quality control was performed for all the four urinary biomarkers data (http://biobank.ctsu.ox.ac.uk/crystal/docs/urine_assay.pdf). Baseline characteristics of UKB participants who had included in the analyses are shown in S1 Table. Details of these measurements can be found in the UKB Data Showcase (http://biobank.ctsu.ox.ac.uk/crystal/). We used rank-based inverse normal transformed phenotype levels in all continuous association tests performed.

### Genotypes

Genotyping was performed with the UK BiLEVE and UK Biobank Axiom arrays (Affymetrix Research Services Laboratory, Santa Clara, California, USA). Initial quality control (QC) was conducted centrally by the UKB, and has been described in detail by Bycroft et al (19). At the time of this study, genotype data was available for 488,377 participants at 805,426 markers; genotype imputation was also conducted centrally using IMPUTE2 and a reference panel from the Haplotype Reference Consortium (HRC) (19). In our analysis, we used the July 2017 release of the imputed genetic marker data, and excluded genetic markers with minor allele count ≤ 30 and imputation quality < 0.8. As a result, the total number of genetic markers included in our analysis was 15,640,977.

### Genetic association analysis

We excluded individuals who had withdrawn consent at the time of this study, who were related, and who did not self-report as white or did not cluster with Europeans based on principal component analysis of genetic data (N=337,542). Unrelated individuals were defined as those who were no closer than the third-degree based on pairwise kinship coefficients by Bycroft et al (19).

In remaining participants, we performed GWAS to identify genetic variants associated with each of the 4 biomarkers (creatinine, N = 327,857 [3.0% missing values]; microalbumin, N = 326,441 [3.3% missing values]; potassium, N = 327,147 [3.1% missing values]; sodium, N = 327,162 [3.1% missing values]). Urine creatinine (μmol/L; ID 30510), potassium (mmol/L; ID 30520) and sodium (mmol/L; ID 30530) were analyzed as continuous traits, while microalbumin (mg/L; ID 30500) was dichotomized into a binary trait (≤30 mg/L = 0 and >30 mg/L = 1)(20). The rationale for dichotomizing urinary microalbumin was that the distribution was bimodal, as expected in a community-based sample where the majority has no measurable urine albumin. We conducted association analyses with PLINK (version 2.0) (21) using linear or logistic regression of biomarker levels on imputed genotypes assuming an additive model between phenotypes and genotype dosages. We adjusted all models for age, sex, batch (3 levels; UK BiLEVE, UK Biobank release 1 and UK Biobank release 2) and the first ten genotype principal components, and restricted association analyses to single nucleotide polymorphisms (SNPs) with minor allele count ≥ 30 and imputation quality information score (info) ≥ 0.8.

We pruned results based on distance retaining variants with the strongest associations within this distance for downstream analysis. We identified regions containing one or more SNPs with p<5e-8 (“index SNPs”) by screening a window of 500kb adjacent to the first index SNP on each chromosome. If no additional SNPs were identified, the region was limited to that specific SNP, and screening was continued at the next index SNP. If additional index SNPs were present in this 500kb window, the window was expanded by a distance of 300kb from the last SNP in search of additional SNPs with p<5e-8 until no more SNPs with p<5e-8 within the next 300kb could be found. From this pruning, we identified a number of regions containing one to several index SNPs and assigned the SNP with lowest p-value within each region as the lead SNP. Within each region, we repeated association analysis after including all index SNPs in the region as well as lead SNPs from other regions on the same chromosome. We considered any SNP with a p<5e-8 in this final analysis to be an independent locus. Regional plots were created for the association test results at significant loci using LocusZoom v1.4 (22).

### LD-score regression

We applied LD-score regression (23) using available GWAS summary statistics from the LD Hub database (24) to evaluate genome-wide heritability (h^2^_g_) of the four urine biomarkers and to identify genetic correlation with other traits. We used pre-calculated European LD scores and restricted the analysis to SNPs found in HapMap Phase 3 (25). We evaluated pairwise genetic correlations of all four urine biomarkers, and between each biomarker and a total of 27 other traits. Specifically, we analyzed correlations with anthropometric (body mass index [BMI], body fat percentage, height, obesity, waist-hip ratio [WHR]), cardiovascular (atrial fibrillation, coronary artery disease, diastolic and systolic blood pressure, heart rate, ischemic stroke), glycemic (T2D, fasting glucose and insulin, 2-hour glucose adjusted for BMI, and glycated hemoglobin), lipid (high and low density lipoprotein cholesterol [HDL-C and LDL-C], triglycerides and total cholesterol), hematologic (mean platelet, platelet count) and kidney (CKD, estimated glomerular filtration rate of creatinine [eGFRcrea] and cystatin c [eGFRcys]) traits. A conservative Bonferroni-corrected threshold of 4.46e-04 (adjusting for 28 traits ^*^ 4 biomarkers) was used to identify significant correlations.

### Functional analysis

We evaluated all genome-wide significant loci for functional impact using variant annotations, gene-level analysis, and colocalization analyses. We annotated chromatin states of all significant SNPs based on the 15-state model used in the NIH Roadmap Epigenomics study (26), and added functional annotations using RegulomeDB categories (27). We used predicted gene functions as implemented in DEPICT (28) to provide biological interpretation of association signals and identify relevant tissues contributing to the signals. To balance power with specificity, DEPICT analysis was performed in two ways: 1) by including independent loci at genome-wide significance; 2) by including loci at a lower significance level (p ≤ 10^−5^) in the GWAS after pruning variants in high LD (r^2^ > 0.05). Apart from a SNP-level GWAS, we conducted a MAGMA gene-level association analysis using gene-level tests as implemented in the FUMA GWAS platform (29). We explored pathways and tissues that were significantly enriched for associated genes using MAGMA (30). Tissue specificity was determined in 53 tissues based on gene expression data from the Genotype Tissue Expression project (GTEx V6) (31). We also determined whether the genes identified by gene-level associations were overrepresented in differentially expressed gene sets (DEG) for each tissue as implemented in FUMA. In these gene-based tests, two-sided Student’s t-tests were performed per gene per tissue against all other tissues. After Bonferroni correction, genes with corrected P-value <0.05 and absolute log fold change ≥ 0.58 were defined as a DEG set in a given tissue. We distinguished between genes that are upregulated and downregulated in a specific tissue compared to other tissues, by taking the sign of t-score into account. Genes were tested against those DEG sets by hypergeometric tests to evaluate if the prioritized genes are overrepresented in DEG sets in specific tissue types.

### Colocalization with eQTL and mQTL summary statistics

We performed summary data-based Mendelian Randomization (SMR) (32) with blood cis-eQTL and cis-mQTL data to evaluate the evidence for colocalization between biomarker GWAS and white blood cell gene expression or methylation signals. We used colocalization to refer to evidence of a causal mechanism between expression and/or methylation and the GWAS signal. eQTL and mQTL colocalization analyses were performed at the level of individual probes; eQTL summary statistics were obtained from Westra et al. (33), and mQTL summary statistics were obtained from McRae et al. (34). We tested 5,959 and 93,220 probes which had genome wide significant cis-eQTLs and cis-mQTLs, respectively (QTL p-value < 5e-08), and controlled false discoveries using Benjamini-Yekutieli (5% FDR) control in both analyses. As some genes have multiple gene expression probes, we estimated pairwise LD between all significant eQTL colocalization lead SNPs while analyzing results to ensure that the same colocalization was not reported using multiple probes. Similarly, multiple methylation probes map to a given region; hence, we repeated the above process while reporting mQTL colocalizations as well.

## Acknowledgements

This research has been conducted using the UK Biobank Resource under Application Number 13721. This research was performed with support from National Institutes of Health (1R01HL135313-01).

## Supporting Information

### Tables

**ST1 Table. Baseline characteristics of UK Biobank participants who had included in the analyses**.

Abbreviations: SD, standard deviation; T2D, type 2 diabetes; SBP, systolic blood pressure; DBP, diastolic blood pressure; BMI, body mass index; WHR, waist-to-hip ratio.

**S2 Table. GWAS catalog reference for genetic loci (500 bp) of the 78 novel lead variants associated with urine biomarkers in the genome-wide association studies**. GWAS hits are highlighted and associations are sorted by p-value. GWAS catalog references were selected applying the following filters: minimum p values of 5e10-8, minimum samples size of 10000 for continuous traits and 5000 cases for binary traits.

Abbreviations: SNP, single-nucleotide polymorphism; POS, position; CHR, chromosome; PMID, pub med identification number.

**S3 Table. P-value per gene-based genome-wide analysis for urinary biomarkers**.

Abbreviations: CHR, chromosome; SNP, single-nucleotide polymorphism; P, p-value.

**S4 Table. Genetic correlation between urinary biomarkers and other traits**.

Abbreviations: rg, genetic correlation; P, p-value; BMI, body mass index; WHR, waist-hip ratio; AF, atrial fibrillation; CAD, coronary artery disease; DBP, diastolic blood pressure; SBP, systolic blood pressure; HR, heart rate; IS, ischemic stroke; T2D, type 2 diabetes; FG, fasting glucose; FI, fasting insulin; 2hrGluAdjBMI, 2 hours glucose adjusted for BMI; HbA1C, glycated hemoglobin; HDL, high density lipoprotein; LDL, low density lipoprotein; TG, triglycerides; TC, total cholesterol; CKD, chronic kidney disease; eGFRcrea, estimated glomerular filtration rate of creatinine; eGFRcys, estimated glomerular filtration rate of cystatin c.

**S5 Table. P-value of tissue enrichment for urinary biomarkers**.

**S6 Table. Significant expression quantitative trait loci (eQTL) probe colocalizations for creatinine**.

Abbreviations: Chr, chromosome; bp, base position; SNP, single-nucleotide polymorphism; A1, effect allele; A2, other allele; Freq, minor allele frequency; b, beta; se, standard error; p, p-value; GWAS, genome-wide association study; eQTL, expression quantitative trait loci; SMR, summary-based-results Mendelian Randomisation; HEIDI, heterogeneity in dependent instruments.

**S7 Table. Significant methylation quantitative trait loci (mQTL) probe colocalizations for creatinine, potassium and sodium**.

Abbreviations: Chr, chromosome; bp, base position; SNP, single-nucleotide polymorphism; A1, effect allele; A2, other allele; Freq, minor allele frequency; b, beta; se, standard error; p, p-value; GWAS, genome-wide association study; eQTL, expression quantitative trait loci; SMR, summary-based-results Mendelian Randomisation; HEIDI, heterogeneity in dependent instruments; p.BY, Benjamini Yekutieli adjusted p-value.

### Figures

**S1 Figure. Regional association and linkage disequilibrium plots for 23 genome-wide significant loci for creatinine**. The y axis represents the negative logarithm (base 10) of the SNP P value and the x axis represents the position on the chromosome, with the name and location of genes in the UCSC Genome Browser shown in the bottom panel. The SNP with the lowest P value in the region is marked by a purple diamond. The colors of the other SNPs indicate the r2 of these SNPs with the lead SNP. Plots were generated with LocusZoom.

**S2 Figure. Regional association and linkage disequilibrium plots for 20 genome-wide significant loci for microalbumin**. The y axis represents the negative logarithm (base 10) of the SNP P value and the x axis represents the position on the chromosome, with the name and location of genes in the UCSC Genome Browser shown in the bottom panel. The SNP with the lowest P value in the region is marked by a purple diamond. The colors of the other SNPs indicate the r2 of these SNPs with the lead SNP. Plots were generated with LocusZoom.

**S3 Figure. Regional association and linkage disequilibrium plots for 12 genome-wide significant loci for potassium**. The y axis represents the negative logarithm (base 10) of the SNP P value and the x axis represents the position on the chromosome, with the name and location of genes in the UCSC Genome Browser shown in the bottom panel. The SNP with the lowest P value in the region is marked by a purple diamond. The colors of the other SNPs indicate the r2 of these SNPs with the lead SNP. Plots were generated with LocusZoom.

**S4 Figure. Regional association and linkage disequilibrium plots for 38 genome-wide significant loci for sodium**. The y axis represents the negative logarithm (base 10) of the SNP P value and the x axis represents the position on the chromosome, with the name and location of genes in the UCSC Genome Browser shown in the bottom panel. The SNP with the lowest P value in the region is marked by a purple diamond. The colors of the other SNPs indicate the r2 of these SNPs with the lead SNP. Plots were generated with LocusZoom.

**S5 Figure. Q-Q plot for genetic associations for urinary creatinine (a), microalbumin (b), potassium (c) and sodium (d)**.

**S6 Figure. Gene-based genome-wide analysis for urinary creatinine (a), microalbumin (b), potassium (c) and sodium (d)**. The significant genes for each chromosome labeled. Negative log10-transformed P values for each gene (y axis) are plotted by chromosomal position (x axis). The gray line represents the thresholds for genome-wide statistically significant associations (p = 5e-08).

**S7 Figure. Functional categories for the genome-wide significant SNPs for urinary creatinine (a), microalbumin (b), potassium (c) and sodium (d)**.

**S8 Figure. The Regulome database score for the genome-wide significant SNPs for urinary creatinine (a), microalbumin (b), potassium (c) and sodium (d)**.

**S9 Figure. The minimum (most active) chromatin state for the genome-wide significant SNPs for urinary creatinine (a), microalbumin (b), potassium (c) and sodium (d)**.

**S10 Figure. Tissue enrichment for urinary creatinine (a), microalbumin (b), potassium (c) and sodium (d)**.

**S11 Figure. Differentially Expressed Gene (DEG) Sets across 30 general tissue types for urinary creatinine (a), microalbumin (b), potassium (c) and sodium (d)**. Significant enrichment at Bonferroni corrected P-value ≤ 0.05 are colored in red.

**S12 Figure. Differentially Expressed Gene (DEG) Sets across 53 specific tissue types for urinary creatinine (a), microalbumin (b), potassium (c) and sodium (d)**. Significant enrichment at Bonferroni corrected P-value ≤ 0.05 are colored in red.

